# Perception of Motion Salience Shapes the Emergence of Collective Motions

**DOI:** 10.1101/2023.12.01.569512

**Authors:** Yandong Xiao, Xiaokang Lei, Zhicheng Zheng, Yalun Xiang, Yang-Yu Liu, Xingguang Peng

## Abstract

As one of the most common and spectacular manifestations of coordinated behavior, collective motion is the spontaneous emergence of the ordered movement in a system consisting of many self-propelled agents, e.g., flocks of birds, schools of fish, herds of animals, and human crowds. Despite extensive studies on collective motions, a systems-level understanding of different motion patterns of collective behaviors is still lacking. This further hinders the adoption of bio-inspired mechanisms for applications of swarm robotics. Here, by leveraging three large bird-flocking datasets, we systematically investigate the emergence of different patterns of collective motions: *mobbing, circling*, and *transit*. We find that flocks with higher maneuverable motions (i.e., *mobbing* and *circling*) prefer to evolve a more nested structure of leader-follower (LF) relations and a clear hierarchy to mitigate the damage of individual freedom to group cohesion. In contrast, flocks with smooth motion (i.e., *transit*) do not display this tactful strategy to organize the group. To explain this empirical finding, we propose a measure based on the perception of motion salience (MS) to quantify the trade-off between individual freedom and group cohesion. Moreover, we perform the correlation analysis between LF and MS, finding that individuals with higher MS tend to lead the group with higher maneuverable motions. Those findings prompt us to develop a swarm model with adaptive MS-based (AMS) interactions and confirm that AMS interactions are responsible for the emergence of nested and hierarchical LF relations in the flocks with highly maneuverable motions. Finally, we implement AMS interactions in swarm robotics that consists of ∼10^2^ miniature mobile robots. The swarm experiments of collective following and collective evacuation demonstrate that AMS interactions not only empower the swarm to promptly respond to the transient perturbation but also strengthen the self-organization of collective motions in terms of temporal cognition.

## INTRODUCTION

The collective motion of organisms, e.g., flocks of starlings, colonies of army ants, schools of barracudas milling, and herds of zebra, is one of the most pervasive and spectacular manifestations of coordinated behavior. Collective motion has been studied for decades from biological, physical, and engineering perspectives^1,2,3,4,5,6,7,8^. The bird flocking is one of the most extensively studied examples of collective motion. For example, the studies on starlings^9,10,11^, pigeons^12^, jackdaws^13,14,15^, and chimney swifts^16^ proposed the topological^10,14^, metric^16^ or plastic^15^ interactions in the flocking models to explain different motion patterns of collective behaviors.

Despite the explanatory success at the macro-behavioral level, the proposed topological, metric, or plastic interactions are *phenomenological* in essence. *First*, most flocking models circumvent the key factor of individual perception at the beginning of modeling. For example, they do not explicitly encode the perception stream to process the sensory information perceived from neighbors’ relative motions but rather feed neighbors’ movement information directly to the flocking models^1^. Beyond these phenomenological descriptions of local interactions, more mechanistic models have advocated relating the visual projection and motion response to elicit vision-mediated movement coordination^17,18,19,20^. However, these models filter out the context of motion cues due to the linear or Boolean-like visual projection as a kind of transient perception result, which is problematic for flocks with highly maneuverable motions. *Second*, the existing flocking models always ignore discovering the intrinsic relations between group organization and individuals’ motion characteristics, which underlies the emergence of self-organization in the formation process of collective motions. It is because these relations not only bridge perceptions and decisions (through the combination of local interactions) at the individual level, but also answer the question about what kind of motion characteristics related to leader-follower relations in shaping the emergence of collective motions at the group level. *Third*, existing phenomenological or visual projection models haven’t facilitated the adoption of bio-inspired mechanisms for the applications of swarm robotics^21,22^. Indeed, most of the theoretical models prescribe the position-based and average interactions to yield simple alignment, attraction, and repulsion^23,24,25,26^, while real applications require more sophisticated and highly maneuverable behaviors, e.g., collective anti-predators^15^, collective turn^27^, cooperative transportation^28^ or excavation^29^, collective chase or escape^30,31,32^, etc. Obviously, for accommodating different kinds of collective tasks, designing bio-inspired swarm robotics is inextricably linked to revealing the fundamental mechanisms underlying different motion patterns of collective behaviors from real flocks.

A systems-level understanding of collective motions requires us to integrate a comprehensive research chain from biological observation to bionic mechanism to bio-inspired swarm robotics. To achieve that, in this work, we analyzed high-resolution movement data of bird flocks with different motion patterns, i.e., mobbing, circling, and transit flocks^13,14,15,16^. Leveraging recent advances in the leader-follower (LF) relationship analysis^12^ and the structure analysis of complex systems^33-37^, we aimed to unveil the motion characteristic needed by an individual to lead the flock.

*First*, we found that flocks with higher maneuverable motions (e.g., mobbing and circling) prefer to have a more nested structure of LF relations and a clear hierarchy to maintain group cohesion. *Second*, we proposed a measure based on the perception of motion salience (MS) to quantify the trade-off between individual freedom and group cohesion and demonstrated that individuals with higher MS tend to lead the group with higher maneuverable motions. *Third*, we modeled leaders’ characteristic motions as an adaptive MS-based (AMS) interaction rule about how the focal agent aligns with its neighbors after perceiving their motions. *Finally*, we adopted the AMS interaction rule learned from real flocks for swarm robotics consisting of ∼10^2^ two-wheel differential mobile robots. Through extensive swarm experiments of collective following and collective evacuation, we found compelling evidence that AMS interactions facilitate the emergence of collective motions for swarm robotics.

## RESULTS

### Empirical data analysis

To reveal the intrinsic interactions in collective motions, we leveraged the high-resolution movement data of bird flocks with different motion patterns (i.e., *mobbing, circling* and *transit*)^13-15,16^. The mobbing and transit datasets were obtained from the video recording of 3D movements of all individuals within flocks of wild jackdaws (*Corvus monedul*) in Cornwall, UK^13-15^. The circling dataset contained 3D tracks reconstructed from video recordings of a flock of ∼ 1,800 chimney swifts (*Chaetura pelagica*) in the field when the swifts entered an overnight roost in Raleigh, USA^16^. The mobbing flocks recorded the collective anti-predator events during which individuals gathered together to inspect and repel a predator (**Fig. 1a** and Supplementary Video 1). The circling flocks with hundreds of chimney swifts displayed the circling approach pattern from surrounding areas near a roost site (**Fig. 1b** and Supplementary Video 2). The transit dataset showed the highly ordered and smooth movement of jackdaw flocks flying towards their winter roosts (**Fig. 1c** and Supplementary Video 3). See Supplementary Sec.1 and Supplementary Fig. 1 for detailed information about the data collection and processing of three datasets. In this work, we totally got 140, 94, and 1483 tracks of mobbing, circling, and transit flocks, respectively. The three datasets could be categorized into two kinds of motion patterns: the first two (*mobbing* and *circling*) displayed highly maneuverable motion, i.e., collective sharp turn to drive away predator or collective circling near a roost site, while the last one (*transit*) is much smoother and highly ordered to move towards the winter roosts (see Supplementary Sec.3 and Supplementary Figs.2-10 for the overview of flocking trajectories).

**Figure 1.**
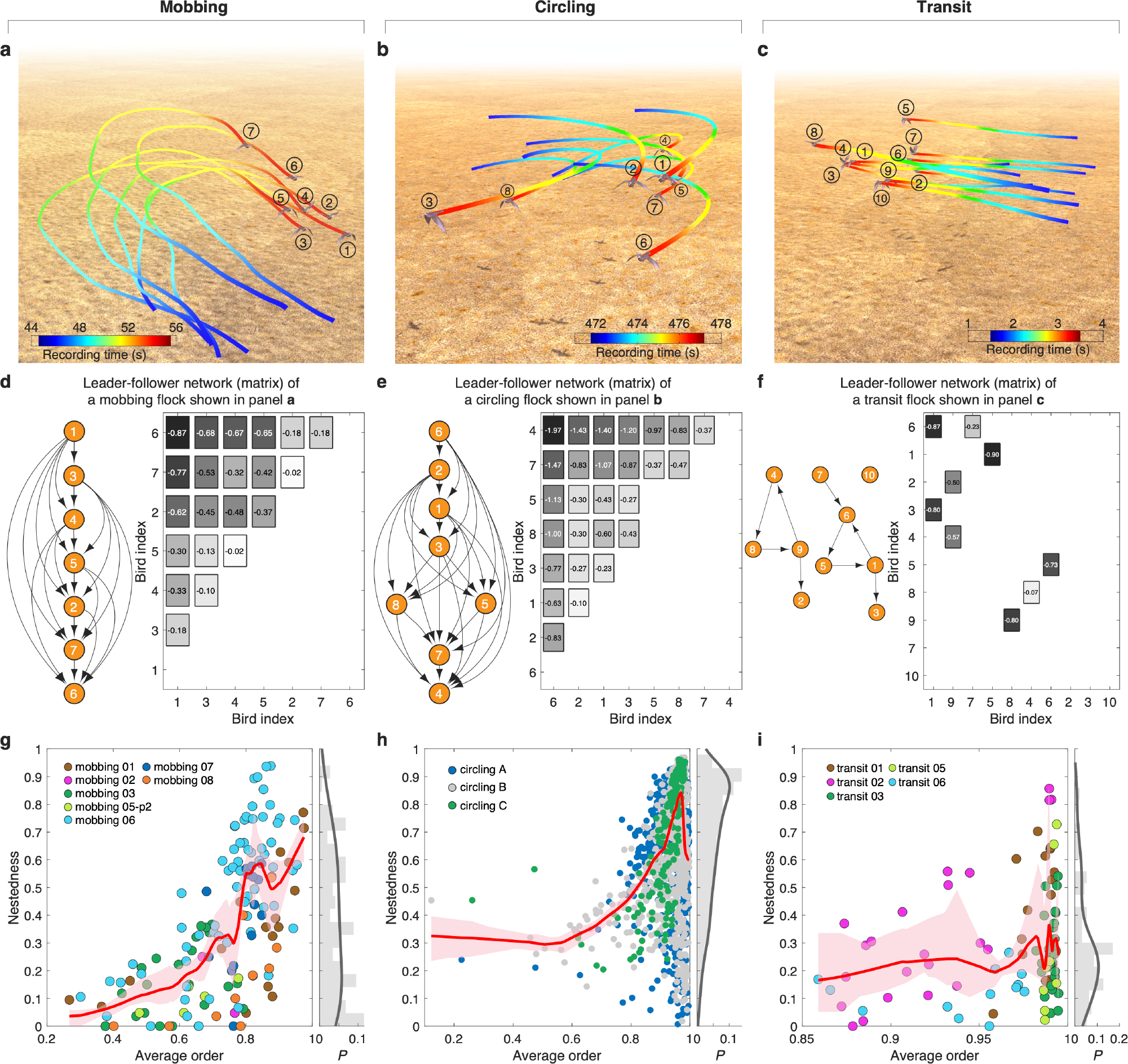
The leader-follower relations of different patterns of collective motions. The flocking trajectories are from three datasets: mobbing (**a**), circling (**b**) and transit (**c**), which are categorized into maneuverable (mobbing and circling) and smooth (transit) modes of collective motions. In panels a-c, the gradient color from blue to red corresponds to the recording time from beginning to end. **d-f**, The LF relation matrix and corresponding LF network of three flocks shown in panels a-c, respectively. The negative values 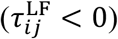 in LF relation matrix represent that individual-*j* leads individual-*i*, and equals the directed edges pointing from leader-*j* to follower-*i* in the corresponding LF networks. The box colors from black to white correspond to the descending order of absolute value of 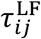. The nestedness of LF networks as a function of average order for three flocking datasets of mobbing (**g**), circling (**h**) and transit (**i**). The right marginal plots show the nestedness distribution of LF networks. The order parameter is a temporalmeasurement evaluating the motion polarization of a flock at time *t*, i.e., 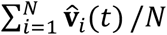, and the average order is the mean of polarization over the whole time. In panels d-i, the LF relation matrices are calculated from the whole period of each flock, and the nestedness is measured by NODF. In panels g-i, each point represents a flock from three datasets, and the nonparametric regression and bootstrap sampling are performed to calculate the trend (red curve) and its 94% confidence interval (red shadow) between nestedness and average order.

#### Flocks with higher maneuverable motions display stronger nested leader-follower relations

To reveal the subtle mechanism of group organization embedded in flocks with different patterns of collective motions, we investigated how LF relations are organized in the flocks. Consider a pair of birds *i* and *j* in a flock within the time period [*t*_*s*_, *t*_*e*_], and the normalized temporal flying direction denoted as 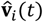 and 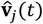, respectively. For time *t* and time interval *τ*, the degree of motion alignment of this bird pair is defined as 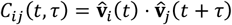. Here the sign of *τ* means the lag (*τ* < 0) or advance (*τ* > 0) time of focal *i* from the view of neighbor *j*. For a given *τ*, we averaged *C*_*ij*_(*t, τ*) over all the time stamps *t* from the period [*t*_*s*_, *t*_*e*_], yielding ⟨*C*_*ij*_⟩(*τ*) =⟨*C*_*ij*_(*t, τ*) ⟩, which indicates the LF relation between birds *i* and *j* in the flock within [*t*_*s*_, *t*_*e*_] (see **Methods**)^12^. The value of 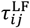 that makes ⟨*C*_*ij*_⟩ (*τ*) reach the maximal value is referred to as the leading or lag time of this pair’s LF relation, i.e., the time that individual-*j* could align with individual-*i* with the maximal consensus within the period 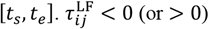 means that the flying direction of individual-*i* lags 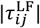seconds behind (or advances 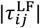 seconds before) that of individual-*j* to keep the maximal consensus, which could be interpreted as a case of individual-*i* following (or leading) individual-*j*, respectively. For example, in a mobbing flock shown in **Fig. 1a**, due to 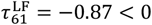, bird-1 is the leader of bird-6, because bird-6 falls 0.87 seconds behind to maintain the maximal alignment with bird-1 (**Fig. 1d** and Supplementary Fig. 12). For a flock within [*t*_*s*_, *t*_*e*_], we constructed an LF relation matrix **T**_LF_(*t*_*s*_, *t*_*e*_) by assigning its (*i, j*)-entry as 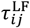if 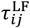 and 0 otherwise (see matrices in **Fig. 1d-f**). An LF relation matrix can also be represented as a directed network, where the directed edge points from the leader *j* to its follower *i* if there exists 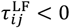 (see LF networks in **Fig. 1d-f**).

Interestingly, we found that the LF network of the mobbing (or circling) flock displays a highly *nested* structure^33-36^ and is completely *hierarchical* (**Fig. 1d**,**e**), which is consistent with previous findings in pigeon flocks^12^. Here, the nested structure means that for those birds with few leaders (i.e., they have a high position in the hierarchy), their leaders also tend to be the leaders of those birds who have many leaders and a low position in the hierarchy. A completely hierarchical structure means that birds with lower positions will never be leaders of birds with higher positions in the hierarchy. In other words, the LF network is a directed acyclic graph (DAG) that has no directed cycles. The hierarchy of a DAG is also known as its topological ordering. However, for the transit flock shown in **Fig. 1c**, its LF relation matrix doesn’t display neither a nested nor a hierarchical structure (**Fig. 1f**).

To systematically study the hierarchy in the flocks with different patterns of collective motions, we calculated the nestedness of LF networks for the three flocking datasets (see Supplementary Sec.4 for details). The distributions of nestedness values displayed in marginal plots of **Fig. 1g-i** demonstrate that the nestedness of transit flocks is significantly lower than that of mobbing and circling flocks (*p*-value = 0.017 and 5.54e-52, respectively, Mann–Whitney U-test). Furthermore, we plotted the nestedness of the LF network of a flock as a function of its *average order* for three flocking datasets. Here, the *order* of a flock at time *t* is a temporal measurement evaluating its motion polarization, and the *average order* of a flock is the average of its order over the entire period of measurement. We found that both mobbing and circling flocks show an overall positive correlation between their average order and the nestedness of their LF networks (**Fig. 1g**,**h**), while this positive correlation is not observed for transit flocks (**Fig. 1i**).

#### Perception of motion salience quantifies the trade-off between individual freedom and group cohesion in collective motions

Overall, the empirical data analysis demonstrates that flocks with highly maneuverable motions tend to have a highly nested structure of LF relations and a clear hierarchy to effectively maintain the group order. This finding prompts us to ask a fundamental question: *What are the interaction rules responsible for the emergence of nested and hierarchical LF relations?* To address this question, we first studied the individual perception of their neighbors’ motion changes. In cognitive science, a seminal model, namely the salience model, and its variants reported the human shifts of visual attention in the spatial and temporal aspects^38,39^. Coincidentally, the empirical evidence in fish schooling revealed the selective attention phenomena that the individual guides its directional decisions according to neighbors’ relative strength of visual feature^40,41,42^, and a large-scale motion-capture system uncovered the visual attention of freely-behaving pigeons^43^. Inspired by these findings, we realized that the individual perception should encode the attention information for the neighbors’ movement changes within a period of time. Therefore, we proposed the individual perception of motion salience (MS) to measure the relative movement changes of neighbor-*j* from the focal individual-*i*’s perception within the period [*t* − *τ, t*], denoted as *M*_*ij*_(*t, τ*). Here *τ*, being 0 < *τ* < *t*, means the lag time for individuals *i* and *j* at time *t*, and is also referred to as the perceiving time. The larger *M*_*ij*_(*t, τ*), the more pronounced movement of individual-*j* felt by individual-*i* during [*t* − *τ, t*] (see **Fig. 2a** and **Methods** for the detailed definition of *M*_*ij*_(*t, τ*)). We incorporated an anisotropic factor α ≥ 0 in the definition of MS to model the fact of forward-oriented preference in biological perception^16,44,45^ (**Fig. 2b**). With increasing α, the ability of individual perceiving movements of its neighbors gradually narrows to the front vision. We emphasized that the definition of MS in this work aims to measure and compare the relative motion changes of multiple neighbors from the context of collective motion cues, which is different from visual selection/search^38,39^ or salient motion detection^46^ in the fields of cognitive science or computer vision.

**Figure 2.**
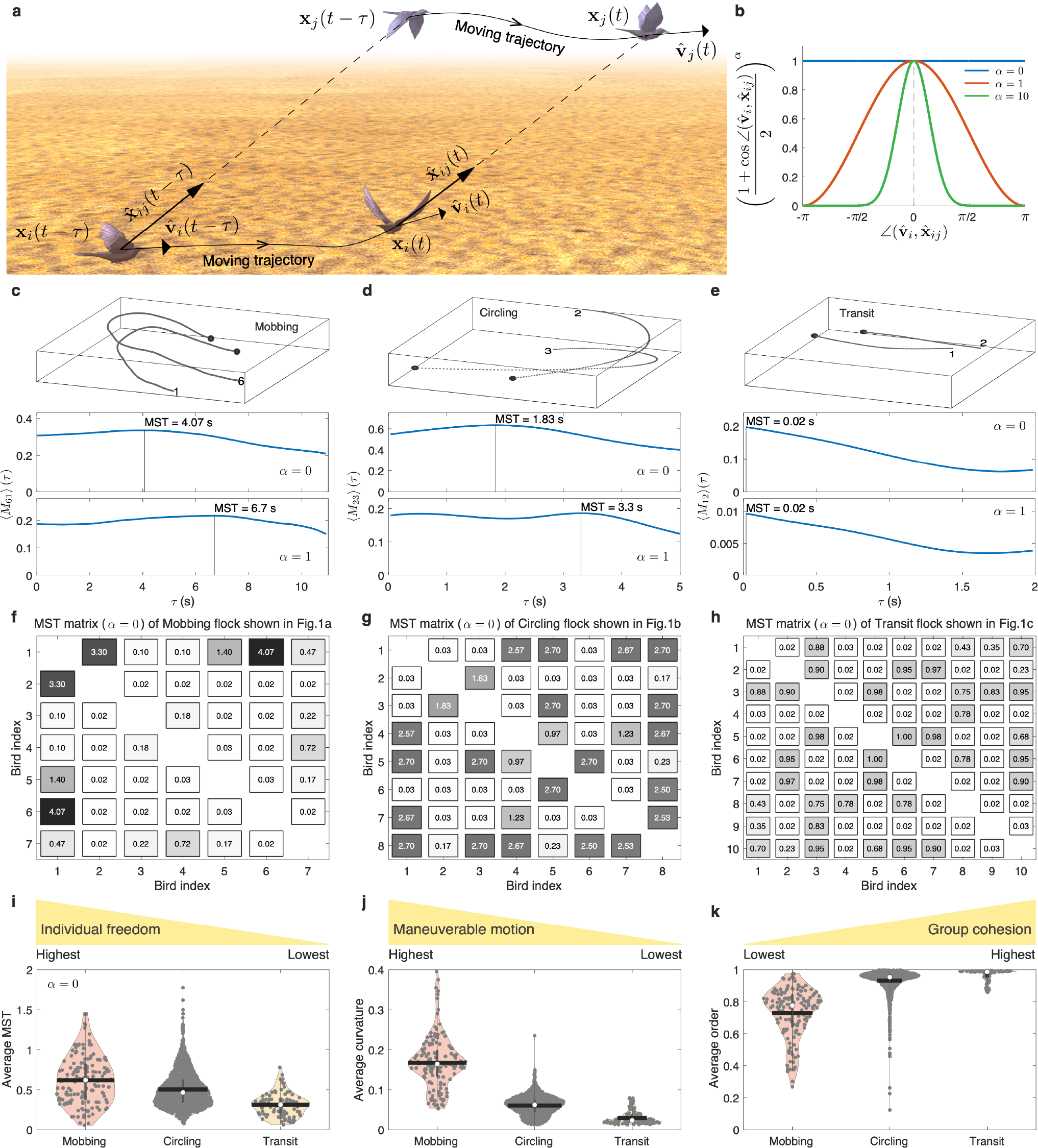
Perception of motion salience in collective motions. **a**, Diagram of MS to quantify the relative movement changes between an individual pair. **b**, The anisotropic effect of motion perception in Eq.(2) mimics that the perceiving ability decays with increasing the sight from front to back. The *x*-axis indicates the heading between two vectors 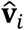and 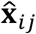, and *y*-axis corresponds to the second or third term of right side of Eq.(2). If α = 0, we ignore the blind area of motion perception. With increasing α > 0, we assumed that the ability of individual perceiving movements of around neighbors gradually narrows to the front vision. ⟨*M*_*ij*_⟩ (*τ*) is the average MS of bird-*i* perceiving bird-*j*’s relative motion changes as a function of *τ* from a flock of mobbing (**c**), circling (**d**) and transit (**e**), respectively. The MST matrix for a flock of mobbing (**f**), circling (**g**) and transit (**h**), respectively. The numbers in the boxes represent MST of each individual pair, and the box colors from black to white maps the descending order of MST. In panels c-h, the flocks used to calculate results are the same with Fig. 1a-c. The distribution of average MST (**i**), average curvature (**j**) and average order (**k**) for three flocking datasets. In panels i-k, each grey dot represents a flock from three datasets, and the white points (or black lines) represent the median (or mean) value.

For a given *τ* > 0, averaging *M*_*ij*_(*t, τ*) over all the time stamps *t* from the period [*t*_*s*_, *t*_*e*_] yields ⟨*M*_*ij*_⟩(*τ*). We defined the maximal motion salience time (MST), denoted as 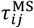, to be the *τ* value where ⟨*M*_*ij*_⟩(*τ*) reaches its maximum. *τ*^*+^ characterizes how long it will take individual-*j* to reach the maximum of relative motion changes from individual-*i*’s perception. If *τ* is shorter than MST, ⟨*M*_*ij*_⟩ (*τ*) could approach to maximum with increasing *τ* firstly. The increment of ⟨*M*_*ij*_⟩ (*τ*) means the motion difference between two individuals could become more and more pronounced. Until *τ* approches to MST, the motion difference reaches the maximum. Beyond MST, two individuals’ movements start to become converged. MST, in essence, equals to the duration of neighbor’s flying freedom. If MST is smaller (or larger), the neighbor has a short (or long) time to move freely relative to the focal one (see **Methods** and Supplementary Fig. 13). **Fig. 2c-e** show the curves of ⟨*M*_*ij*_⟩ (*τ*) and MST of three individual pairs from mobbing, circling and transit flocks displayed in **Fig. 1a-c**, respectively.

For any flock, within a time period [*t*_*s*_, *t*_*e*_]we could construct the MST matrix. **Fig. 2f-h** show the MST matrices for the mobbing, circling, and transit flocks. We found that, overall, the MST values of the two highly maneuverable flocks (mobbing and circling) are much larger than that of transit. This is consistent with the degree of maneuverability observed from the corresponding flocking trajectories. We then took the average of all the elements in MST matrix to evaluate the degree of flying freedom for a flock. As shown in **Fig. 2i**, the mobbing and circling flocks demonstrate significantly higher average MST than that of transit flocks (*p*-value=1.46e-17 and 2.39e-16, respectively, Mann–Whitney U-test). This is accorded with the fact that the mobbing and circling flocks have a higher degree of maneuverability (and a lower degree of average order) than transit flocks (**Fig. 2j**,**k**). Here the degree of maneuverability is quantified by the average curvature of the flocking trajectories. Furthermore, both mobbing and circling flocks display a significant falling trend of average MST with the increment of the average order. Conversely, the transparent negative correlation between average MST and average order is not observed in the majority of transit flocks (Supplementary Fig. 14a). If we consider the anisotropic effect of motion perception (α > 0), the decreasing rank of average MST and the transparent negative correlation between average MST and average order in three flocking datasets are consistent with the case of α = 0 (Supplementary Fig. 14b,c).

The above phenomena in different patterns of collective motions demonstrate that the smaller the average MST, the smaller the individual freedom, and the higher the group order. This implies that the perception of MS could quantify the trade-off between individual freedom and group cohesion in collective motions. For example, for flocks with maneuverable motions, i.e., mobbing and circling, since individuals have higher MST (or more freedom) to loosely couple with neighbors (induces lower group order), the flock could maintain the flexible motion maneuverability in response to external stimuli, i.e., driving away predators or hovering near a roost site. By contrast, the transit flocks with lower MST, emerging the straight flying pattern, require individuals to sacrifice their own freedom to keep a well-ordered alignment with neighbors. Therefore, the findings of MS explain that sacrificing individual freedom makes well-organized LF relations unnecessary for the transit flocks, while flocks with maneuverable motions need to work subtly to offset the disadvantages of individual freedom.

#### Individuals with higher motion salience tend to lead the group in the maneuverable motions

According to Eq.(2), we could construct MS matrix *M*(*t, τ*) = [*M* _*ij*_(*t, τ*)]_*N*×*N*_ and derive the average MS of each individual *M*_*i*_ (*t, τ*) by averaging each column of *M*(*t, τ*). Moreover, besides the nestedness value capturing the hierarchy of LF relations at the group level, we also calculated the local reaching centrality^37^ to quantify the leading tier of each individual in an LF network derived from a flock within the time period [*t* − *τ, t*], labelled as *L*_*i*_(*t, τ*) ∈ [0,1] (see **Methods**). The larger the *L*_*i*_, the higher the leading role of individual-*i*. For example, in the LF network of **Fig. 1d**, bird-1 at the top, leading all the downstream individuals, has the largest *L*_1_ = 1, while *L*_6_ = 0 of bird-6 is the least because it locates at the lowest layer of the LF network. An in-depth study of the correlation between LF and MS enables us to disclose the role of individual motion characters during the group formation to answer a fundamental question: *what kind of motion characteristic does an individual possess to lead the flock?* Leader’s motion character could be interpreted as the interaction rule about how the focal aligns with neighbors after perceiving their motions. For example, according to the mobbing flock shown in **Fig. 1a**, we calculated the LF network and MS matrix (*α* = 0) from a short period [*t* − *τ, t*], and then derived each individual’s leading tier (*L*_i_(*t, τ*)) and average MS (*M*_*i*_ (*t, τ*)) (**Fig. 3a**). Interestingly, the two vectors composed of *L*_*i*_ (*t, τ*) and *M*_*i*_ (*t, τ*) show a highly positive correlation. Finally, for the mobbing flock within [*t* − *τ, t*], we computed the Spearman correlation coefficient (*ρ*) between the two vectors composed of *L*_*i*_ (*t, τ*) or *M*_*i*_ (*t, τ*) over different combinations of *t* and *τ* (see **Fig. 3b1** for *α* = 0, **Fig. 3c1** for *α* = 1 and Supplementary Video 4). Besides, we performed the correlation analysis between LF and MS for another two flocks shown in Fig. 1b,c (**Fig. 3b,c**). Interestingly, if we considered the forward-oriented preference of visual perception (*α* = 1), the two flocks with maneuverable motions (mobbing and circling) display positive correlations between LF and MS for almost all the different combinations of *t* and *τ* (**Fig. 3c1,c2**), meaning that the individual with larger MS tends to play the higher-tier leading role for the vast majority of flying time of the flocks. By contrast, the transit flock displays well-mixed positive and negative correlations between LF and MS at different combinations of *t* and *τ* (**Fig. 3c3**).

**Figure 3.**
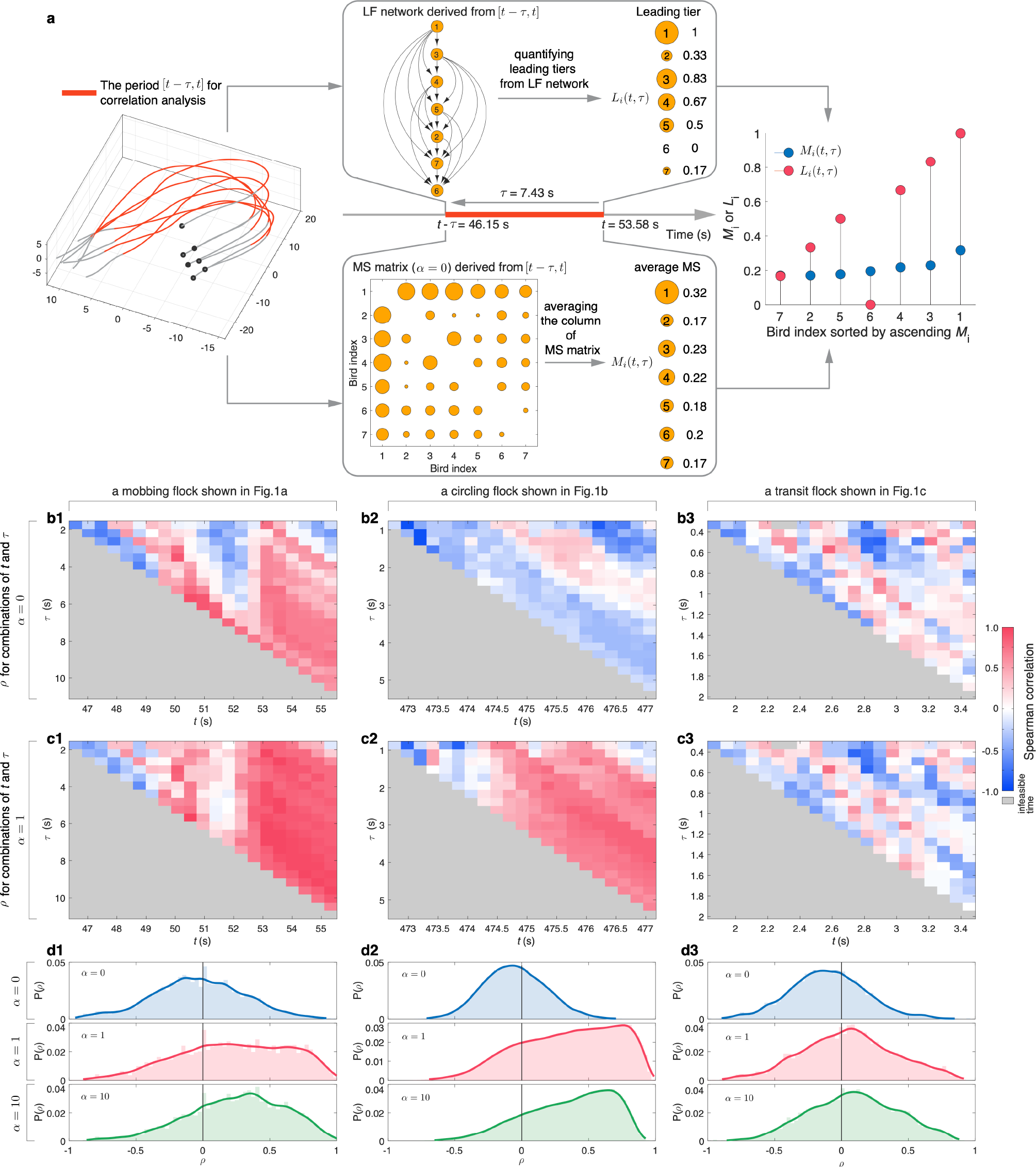
Correlation analysis between LF and MS in collective motions. **a**, For a flock within the period [*t* − *τ, t*] (highlighted by red), first we could construct the LF network and MS matrix (*α* = 0), and consequently derive the leading tier *L*_*i*_ (*t, τ*) and average MS *M*_*i*_(*t, τ*) of each individual. Interestingly, two vectors composed of *L*_*i*_ (*t, τ*) and *M*_*i*_ (*t, τ*) show the highly positive correlation. Finally, we performed the Spearman correlation (*ρ*) between two vectors composed of *L*_*i*_ (*t, τ*) or *M*_*i*_ (*t, τ*) over the different combinations of *t* and *τ*. The heatmap of *ρ* under *α* = 0 (**b**) and *α* = 1 (**c**) for mobbing, circling and transit flocks shown in Fig. 1a-c. The gradient color from blue to white to red maps *ρ* ∈ [−1,1], respectively. The gray color means the infeasible time for Eq.(2), i.e., *t* − *τ* < 0. **d**, The distribution of *ρ* over different combinations of *t* and *τ* from flocking datasets of mobbing, circling and transit when *α* = 0, *α* = 1 and *α* = 10. In panels b-d, for each flock, we took 21 time stamps for *t* and 22 time points for *τ* no matter how long a flock lasts.

Next we extended the correlation analysis to three flocking datasets (**Fig. 3d**). When *α* = 0, three flocking datasets display quite similar *ρ* distribution centered around 0. However, if we consider the forward-oriented sight of birds^16,44,45^ (*α* > 0), the two maneuverable flocking datasets display a *ρ* distribution dominated by positive values, while the transit flock datasets still display a *ρ* distribution centered around 0. The domination of positive *ρ* occurred in the situation of forward-oriented preference of perception (*α* > 0) accords with the spatial structure of the leading position, which aims to investigate the relations between the leading tier and fraction of neighbors present in the front view (Supplementary Sec.5). It demonstrates that in mobbing and circling flocks, the higher leading tier an individual occupies, the more likely it will be at the front of flock to lead the group. On the contrary, the transit flocks do not show any preference between the leading tier and neighbors’ relative position (Supplementary Fig. 16).

### A swarm model with adaptive MS-based interactions

The empirical findings of bird flocks inspire us to build a swarm model with adaptive MS-based (AMS) interactions, i.e., the focal individual adaptively scales its neighbors’ influences through perceiving their MS (**Fig. 4a** and **Methods**). We hypothesized that AMS interactions could be responsible for shaping the emergence of nested and hierarchical LF relations in collective motions. For comparison purposes, we also considered two MS-free interactions: (i) adaptive interaction based on transient heading difference^47,48^ (ATHD), which could adaptively make the neighbors with larger (or smaller) transient heading difference exert larger (or smaller) influences on the focal individual (**Fig. 4b** and **Methods**); (ii) average interaction^23^: the standard Vicsek model that the focal agent equally interacts with its neighbors located in the sensing radius (**Fig. 4c** and **Methods**).

**Figure 4.**
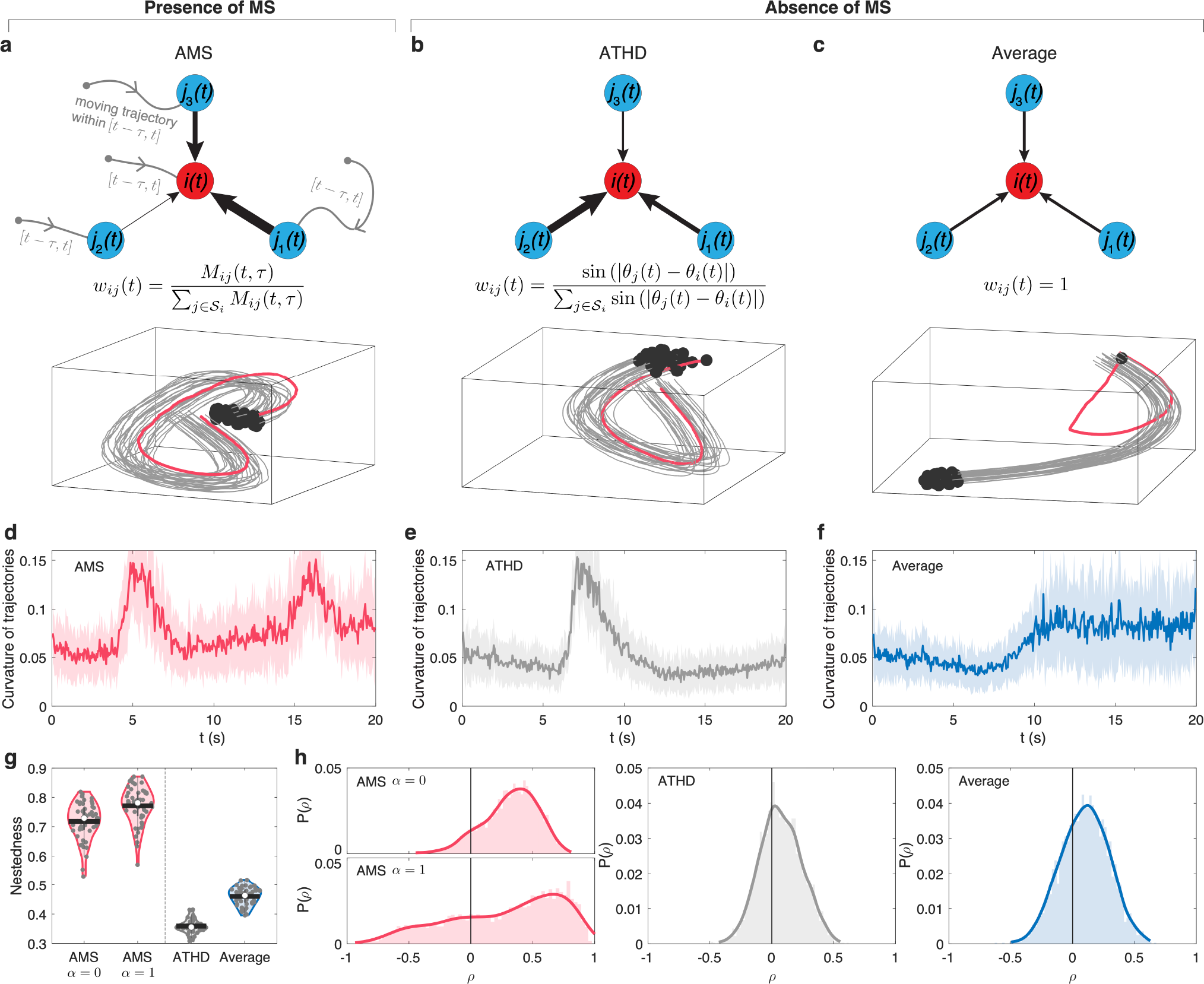
Comparison of self-propelled particle model in presence or absence of MS. We used a self-propelled particle model that particles follow the local interaction rules, i.e., AMS (**a**), ATHD (**b**) and average interaction (**c**). The additional potential well, imposed on one of individuals (red trajectories), aims to lead the flock come back to origin. The swarm size is 30 particles. The black points represent the end of trajectories. In panel a, the flocking trajectories are generated by AMS with *α* = 0. **d-f**, The temporal curvature of flock trajectories respectively shown in panels a-c. In panels d-f, the solid curves represent the average of trajectory curvatures from 30 particles, and the shadow area represents the standard deviation (SD). **g**, The distribution of nestedness of LF networks derived from flocking trajectories using AMS, ATHD and average interaction. The white points (or black lines) represent the median (or mean) value. **h**, The distribution of Spearman correlation (*ρ*) between LF and MS over different combinations of *t* and *τ* from flocking trajectories using AMS, ATHD and average interaction. For each flock we took 10 time stamps for *t* and 11 time points for *τ* to perform the correlation analysis between LF and MS. In panel g and h, we ran 50 independent simulations for each interaction type.

To verify the hypothesis of AMS interactions, we used a self-propelled particle model, where agents follow the local interaction rules, and the additional potential well is imposed on one of individuals to lead the flock come back to the origin (see **Methods**). For example, starting from the same initial conditions, the flock trajectories in **Fig. 4a-c** show that AMS and ATHD interactions could make the informed individual (red trajectories) successfully lead the whole flock, but the average interaction fails. Interestingly, we found that even the adaptive interactions from AMS and ATHD work well to lead the group, AMS interactions could generate more maneuverable motions than that of ATHD interactions within the same period of time (**Fig. 4d,e**). This is because that ATHD interactions consider the adaptive interaction from transient salience and hence constraint the individual freedom. The performance of AMS interactions indicates the advantage of balancing the group cohesion and individual freedom. Furthermore, compared with ATHD and average interactions, AMS interactions yield highly nested LF relations (**Fig. 4g**) and domination of positive correlation between LF and MS (**Fig. 4h**). The results again demonstrate that AMS interactions successfully capture the key characters revealed from real bird flocks.

### Application to swarm robotics

Considering the impact of AMS interactions in the swarm models, we further adopted it to swarm robotics to demonstrate the advantages of self-organization in two swarm experiments. For simplicity in the experiments, we ignored the blind area of motion perception and thus set *α* = 0 in AMS.

#### Collective following experiments

Collective following experiments aim to mimic the ability to respond with agility to rapid changes of moving directions while maintaining the group cohesion, subject to the perturbations from external stimuli or interior neighbors. The inspiration of experimental set-up comes from collective foraging, for instance, an informed individual leads the group to the multiple destinations that frequently changes during foraging^49,50^. In the collective following experiments, we supposed that one of the swarm acts the informed role to move towards the nearest target, and the rest align with neighbors through AMS. To simulate the rapid motion changes for the informed robot, we artificially set 10 targets to guide the moving direction of informed robot through the switch among three different states: living, sleeping and arrived. See Supplementary Sec.6.1 for detailed information about how the informed robot moves according to the target states. Besides, we used *r*(*t*) ∈ [−1,1] to measure the temporal collective response of a flock to rapid motion changes of the informed robot. *r*(*t*) = 1 means the perfect response that the swarm could ideally copy the velocity of informed robot at the moment *t*, while *r*(*t*) = −1 indicates the worst response that the rest of swarm move oppositely to the informed robot. We also used *R* ∈ [0,2] to cumulatively measure the general performance of collective response for a period of time (see Supplementary Sec.6.1). The less *R*, the better collective response.

The experimental results demonstrate that, using AMS interactions, the swarm with 50 robots not only successfully follow the informed robot to reach 10 targets in sequence (**Fig. 5a** and Supplementary Video 5), but also quickly respond to the motion changes of informed robot (**Fig. 5b**). In particular, when the informed robot drastically changes its heading towards the nearest living target, the swarm could speedily recover to the consensus state (see the sudden drops and meteoric rises of *r*(*t*) in **Fig. 5c**). As a comparison, the swarm using average interaction totally fails to follow the informed robot (**Fig. 5d-f** and Supplementary Video 7). **Fig. 5g** demonstrates that when the flock size is single-digit (< 10), there is little difference in *R* for the interaction rule in the presence or absence of MS. However, AMS interactions show a strong advantage in keeping *R* stabilized at a low level (*R* ≈ 0.1 in **Fig. 5g**), no matter what the flock size increases from 10 to 50 robots in real experiments. In addition to the experimental verification of collective following, a simulation platform is developed with the same robot’s physical characteristics. Even if the flock size grows up to 100 in simulations, *R* is still stable at around 0.1 (**Fig. 5h**). Interestingly, **Fig. 5g**,**h** show that the collective response of AMS interactions almost approaches that of ATHD interactions. The latter could be considered as a kind of theoretically ideal response to neighbors’ perturbations as it could adapt the heading difference without delay.

**Figure 5.**
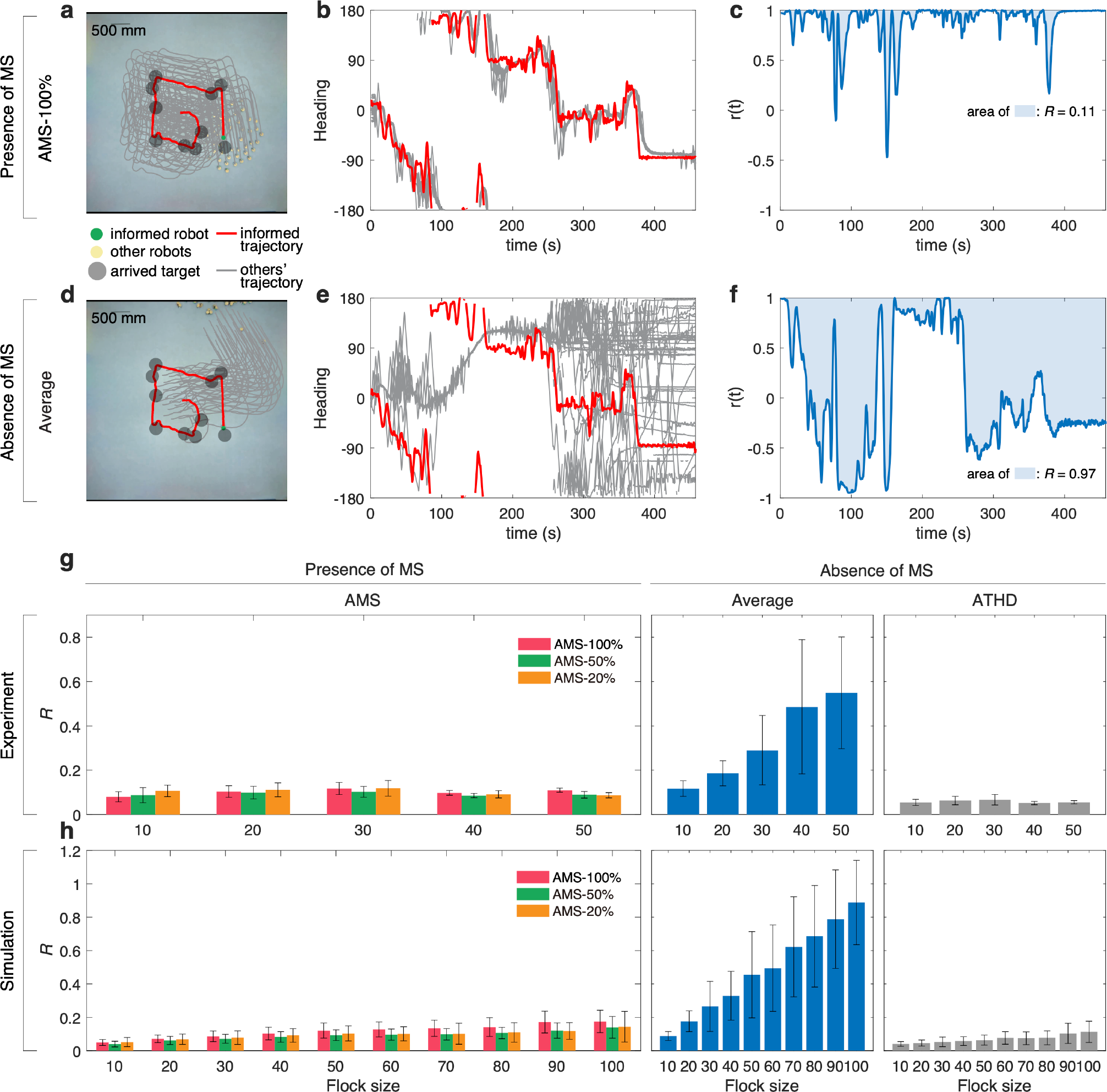
Collective following experiments. **a**, The swarm consisted of 50 robots use AMS to perform the collective following experiment. **b**, Following the informed robots’ movements (red curve), the rest of swarm could promptly respond to stimuli from neighbors and maintain the swarm move together. **c**, The temporal collective response *r*(*t*) for the experment of panel a. AMS makes the cumulative evaluation of collective response *R* approach to about 0.11 (equals to the area of light bule zone). **d-f**, For the same experiment setting of panel a, i.e., the movement of informed robot, the swarm using average interaction could not respond to heading changes of informed robot in time and totally fail to execute collective following. **g**, *R* as a function of swarm size up to 50 robots in experiments for different interaction types. **h**, In the simulations with the same robot’s motion characteristic, *R* as a function of swarm size up to 100 robots for different interaction types. Here AMS-*x*% represents the focal robot only adaptively aligns with those neighbors who cumulatively possess the top *x*% MS. Note that ATHD could be the ideal condition of responding to neighbors’ perturbations because the individual could be able to immediately and adaptively alter the influences from neighbors based on heading difference. In panels g,h, the error bar represents the standard deviation (SD) calculated from 10 independent experiments and 100 independent simulations.

To further reflect the greater attention given to neighbors with higher MS, we introduced a tunable parameter *x*% to AMS interactions: the focal individual only aligns with its neighbors who cumulatively possess the top *x*% MS (see **Methods**). Both the real experiment and simulation results demonstrate that even if just the top 20% or 50% MS is involved, the collective response performs almost the same as that used 100% MS (**Fig. 5g**,**h**). Besides, we systematically investigated the collective response as a function of the top *x*% MS used in AMS interactions. It indicates that the smallest *R* occurs between the top 20% and 40% MS, and *R* slightly increases with the increment of the value of *x*% (Supplementary Fig. 19), which coincidences with the theoretical modeling of selective attention through limiting the cognitive capacity of individuals to maximize the flocking performance^51,52,53^.

#### Collective evacuation experiments

Collective evacuation experiments aim to investigate the effect of AMS to strengthen the self-organization of the swarm to evacuate a narrow gate (**Fig. 6a**). Similar to the application of a drone swarm successfully navigating in confined and cluttered environments, we introduced a state-of-the-art framework^54^ to the collective evacuation experiments, including different kinds of interactions in addition to aligned movement, e.g., collision avoidance against inter-agent or agent-wall, guidance to pass through the exit. Here we only introduced AMS to the alignment part instead of average interaction commonly used in previous works^8,54^ and kept the other terms remained. (see Supplementary Sec.6.2 and Supplementary Fig. 20 for detailed information about the swarm model of collective evacuation). Note that this experiment has no informed robot to explicitly lead the swarm to evacuate the narrow exit.

**Figure 6.**
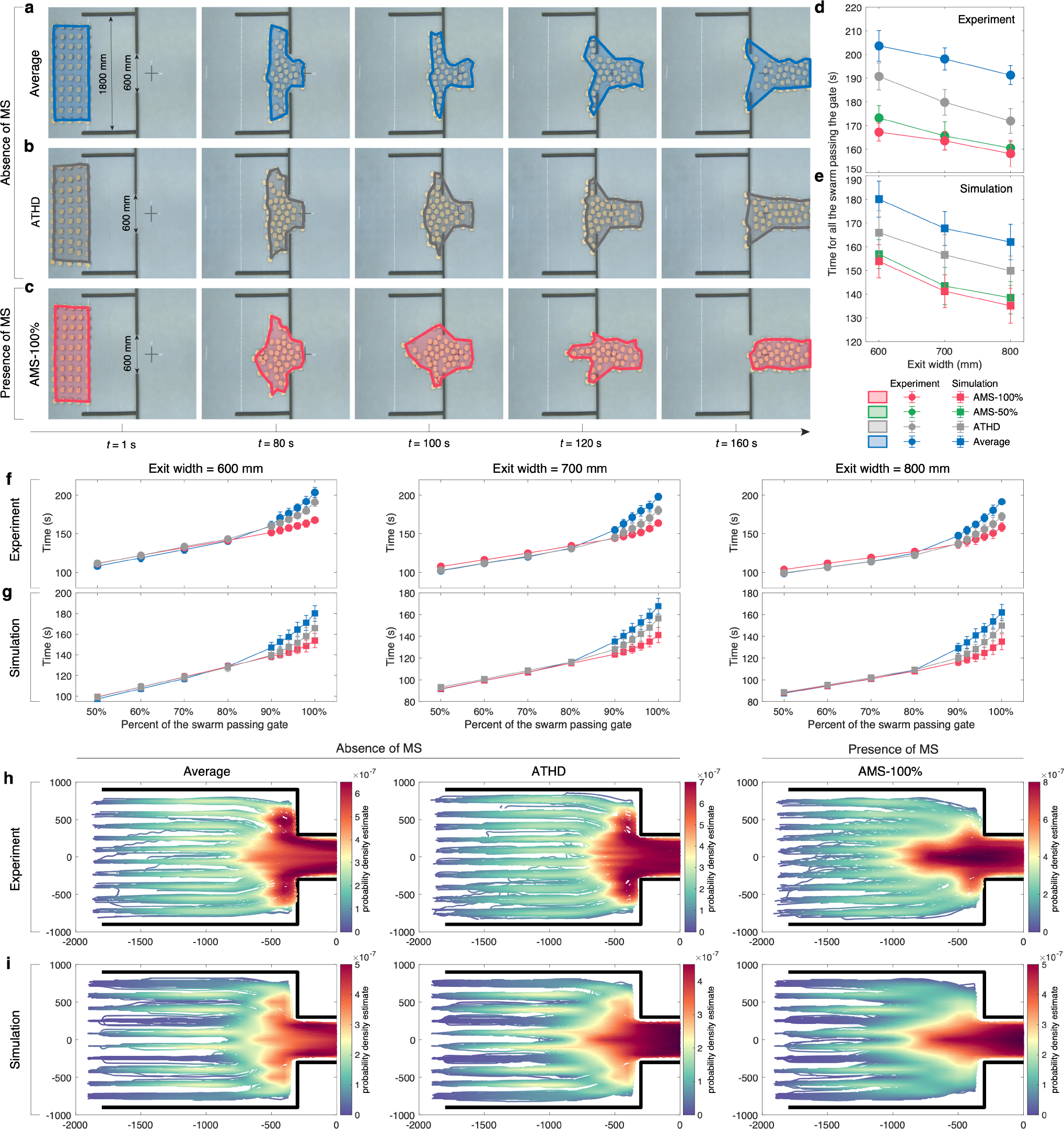
Collective evacuation experiments. The snapshots of collective evacuation experiments for the swarm consisted of 50 robots using average interaction (**a**), ATHD (**b**) and AMS with 100% MS (**c**). The exit width is 600mm. The colored areas highlight the spatial distributions of the swarm. The spending time of all the swarm evacuating the narrow exit as a function of different exit widths from experiments (**d**) and simulations (**e**). The spending time as a function of different percent of the swarm successfully passing the gate from experiments (**f**) and simulations (**g**). In panels d-g, the error bar represents the standard deviation (SD) calculated from 10 independent experiments and 50 independent simulations. The statistics of historical trajectories of all 50 robots in 10 independent experiments (**g**) and 50 independent simulations (**h**). The color scales in panels h,i correspond to the probability density estimate of historical trajectories. The black line indicates the wall. The swarm size is 50 in the experiments and simulations.

The spatiotemporal distributions of the swarm with 50 robots to evacuate the narrow exit (600 mm) are shown in **Fig. 6a-c**. The swarm using average interactions or ATHD interactions matches the common sense that if the exit were not large enough, the swarm would dispersedly occupy the whole inside of the narrow exit^55^ (**Fig. 6a,b** and Supplementary Videos 8-10). Surprisingly, AMS interactions overcome this limitation. A part of the robots at the shoulder of the swarm spontaneously contracts towards the middle when they are far from the exit so that the majority of the swarm could directly face the exit to evacuate (**Fig. 6c**). In particular, we observed that in order to spare space for swarm contraction, some individuals temporarily sacrifice their evacuation time to move towards the opposite direction of the exit (Supplementary Fig. 21). Eventually, the swarm not only evacuate smoothly from the narrow exit as a whole but also cost much less evacuation time than that of average interactions or ATHD interactions.

Therefore, for spending time of swarm evacuation, AMS interactions show tremendous advantages over average interactions and ATHD interactions. For instance, AMS interactions with 100% MS shortens the evacuation time by about 18% compared to the average interaction under three different gate widths (**Fig. 6d,e**). To fully understand the advantage, we recorded the spending time as a function of the percent of the swarm successfully evacuating the gate under different interaction types (**Fig. 6f,g**) and different swarm sizes (Supplementary Fig. 22). For AMS interactions, the evacuation time increases almost linearly with the percent of the swarm that succeeds in exiting. Conversely, the average interaction and ATHD interactions divide the evacuation process into two stages: the first 80% of the swarm spends the time as the same of AMS interactions, but the last 20% evacuate very slowly. The results are consistent with the observed phenomena in the experiments: if using average interaction or ATHD interactions, those robots located directly ahead of the exit could evacuate smoothly, while the robots on either shoulder of the swarm rush straightly towards the side of the exit and consequently induce the congestions. Interestingly, AMS interactions could induce the swarm to spontaneously emerge from the contraction process to directly face the exit and then evacuate as a whole.

The spontaneous emergence of swarm contraction is also validated by the statistics of historical trajectories of all individuals from 10 independent experiments (**Fig. 6h**) and 50 simulations (**Fig. 6i**). (See Supplementary Fig. 23 for statistics of historical trajectories of other exit widths). We noticed that the contraction process evidently emerges only when the individuals perceive the neighbors’ MS for a period of time. For example, if the adaptive influences come from transient differences, such as ATHD interactions, it could improve the performance of swarm evacuation, yet it fails to emerge the swarm contraction (**Fig. 6b** and Supplementary Videos 8-10). Furthermore, different parameters, i.e., swarm size and the top *x*% MS used in AMS interactions, are systematically investigated to verify the generalization of AMS interactions in collective evacuation (Supplementary Fig. 24).

## DISCUSSION

The status quo as it pertains to fundamental mechanisms and potential applications of collective motions could be summarized as: a lack of a comprehensive research chain from biological observation to bionic mechanism to bio-inspired swarm robotics. That has been the case despite numerous studies have been deeply conducted on the above three aspects separately or a part of the research chain. For example, a recent study revealed that group coordination in a flock of sheep results from the information propagation to all group members through a strongly hierarchical, directed interaction network^56^. A study combined the computational and robotic approaches to investigate the strategies for individuals to interact with their neighbors in schooling fish^57^. Swarm of micro flying robots succeeded in navigating and flying through the highly cluttered environments relying on predictive control^54^ or planning/optimization-based methods^58^.

Our work represents a new and substantive departure from the status quo by shifting the focus from one of the specific aspects to a systems-level understanding of collective motions from theory to application. In particular, we modeled the individual perception of MS as an entry point to quantify the movement changes of neighbors. Through the empirical validation on bird flocks, the perception of MS may have major adaptive and evolutional significance to balance the trade-off between individual freedom and group cohesion in collective motions. Since maintaining the flexibility requires more individual freedom to agilely respond to the external perturbations from neighbors or environments, the flocks with highly maneuverable motions (e.g., mobbing and circling flocks) evolve the subtle mechanisms: the nested and hierarchical LF relations could maintain the group order, and the individuals with higher MS prefer to lead the group. Conversely, the highly smoothed flocks do not clearly show the above subtle strategies to organize the group, perhaps due to the less individual freedom. Furthermore, we proposed the MS-based adaptive interaction and adopted this bio-inspired mechanism to swarm robotics. Though we chose the miniature two-wheel differential ground robots as prototypes, the presented swarm experiments of collective following and collective evacuation demonstrated great advantages of AMS in shaping the emergence of collective motions.

It is worth noting that the AMS is very generic for different kinds of collective tasks. *First*, the compelling evidence from two swarm experiments demonstrates that AMS not only empowers the swarm to promptly respond to the transient perturbation but also strengthens the self-organization of collective motions in terms of temporal cognition^59^. For example, collective following experiments indicate that AMS almost approaches the limit of a collective response. Collective evacuation experiments show that AMS makes the swarm spontaneously emerge from the contraction process to directly face the narrow exit and smoothly evacuate as a whole, which effectively avoids the congestion on two sides of the narrow gate. *Second*, AMS is easy to be adopted to different modeling frameworks of collective tasks as it only involves the alignment term. We believe that AMS will have a positive translational impact on the deployment of more advanced autonomous swarm robots^58^ and more sophisticated collective tasks^30,52,60,61^.

Even though the MS-based adaptive interaction paves a promising route to migrate the bionic mechanisms to bio-inspired swarm robotics, we openly admit that it still requires to pay for great efforts, such as precisely and eidetically reconstructing the visual fields of individual perception^30,62,63^, bridging the gap between mathematical modeling of motion salience and perceptual devices on robot^64,65^, etc. On the other hand, recently, neuroscience has made tremendous progress in understanding various aspects of relations between sensory signals and visuomotor stream in larval zebrafish^66,67^. We believe that our findings may also facilitate the design of fully autonomous swarm robotics with artificial neuronal circuits of see-sense-decision-direct, i.e., robots with insect brains^68,69^, from the interdisciplinary exchange between biology, neuroscience, physics, and engineering.

### METHODS

#### Constructing the leader-follower relation matrix of a flock

For an individual pair in a flock with the recording time from *t*_*s*_ to *t*_*e*_ (**Fig. 1a**), we could compute the temporal degree of motion alignment between the focal one’s flying direction 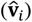 and those of neighbors 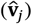 in the form of,

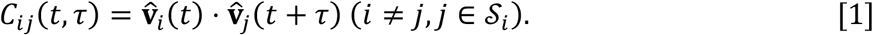

Here *t* is the time stamp, *τ* is the time interval belonging to (*t*_*s* −_ *t*_*e*_, *t*_*s* −_ *t*_*e*_), and 𝒮 _*i*_ indicates the collection of neighbors of the focal *i*. Hereinafter, the vector symbols with hat 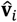means the normalized vector. Note that due to the limitation of recording area in the original observation studies, we roughly assumed that, for a bird flock from mobbing, transit, and circling dataset, 𝒮 _*i*_ represents all individuals appeared in the flock when calculating *C*_*ij*_(*t, τ*).

For a given *τ*, the average of *C*_*ij*_(*t, τ*) over different time stamps within the period [*t*_*s*_, *t*_*e*_] is denoted as ⟨*C*_*ij*_⟩ (*τ*). Thus, we collected the curve of ⟨*C*_*ij*_⟩ (*τ*) as a function of 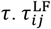 is the value to make the curve of ⟨*C*_*ij*_⟩ (*τ*) reach the maximal value. We disregarded 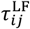 if it locates at first 25% or last 25% of *τ*-axis, as we considered this *τ* to be too short for directional copying^70^. Meanwhile,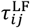 is not considered if the maximal ⟨*C*_*ij*_⟩ (*τ*) is not larger than 0.8. See Supplementary Fig. 12 for details. Note that ⟨*C*_*ij*_⟩ (*τ*) ≠ ⟨*C*_*ij*_⟩ (*τ*) and 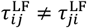, because Eq.(1) does not satisfy the exchange law for an individual pair.

Besides, in the circling datasets, hundreds of swifts hovering over the roost actually contain many sub-flocks due to different spatial locations of individuals in the whole flock. We found that LF relation matrix could successfully classify the circling flocks into sub-communities with more similar motion patterns (see Supplementary Sec.2 and Supplementary Fig. 10 for an example).

#### Quantifying the perception of motion salience from a flock

Motion salience aims to quantify the perception of relative movement changes of neighbor-*j* from the focal *i* within the period [*t* − *τ, t*] in the form of,

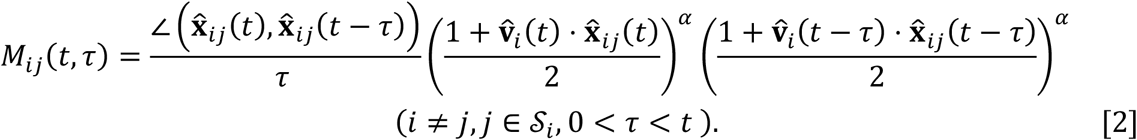

Here 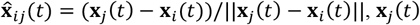 is individual-*j*’s position vector at time *t*, ∠ means the angle between two vectors in 3D Cartesian coordination, and 𝒮 _*i*_ indicates the collection of neighbors of focal-*i*. Similar to Eq.(1), 𝒮 _*i*_ represnets all individuals recorded in the flock when calculating *M*_*ij*_(*t, τ*).

The diagram of definition of MS is shown in **Fig. 2a**. *M*_*ij*_(*t, τ*) comprehensively quantifies the perception of neighbor-*j*’s motion changes relative to the focal individual-*i* involving the position and velocity simultaneously during the period [*t* − *τ, t*]. The last two components in the right side of Eq.(2) considers the anisotropic effect of motion perception, also called the forward-oriented preference of visual perception in birds (**Fig. 2b**). It simulates that the perception ability decays with the sight shifting from the front to back. For example, if 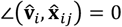, it means that neighbor-*j* locates directly ahead of the focal-*i*’s movement direction, and the focal individual could fully perceive the neighbors because 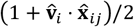 is 1. Once 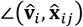 deviates from 0 to π or −π (the horizon axis of **Fig. 2b**), it represents that the neighbor-*j* gradually moves from the front to back of the focal-*i*, and assumes the ability of motion perception could increasingly diminish since 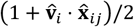 decreases from 1 to 0 (the vertical axis of **Fig. 2b**). Here *α* (only taking *α* ≥ 0) controls the anisotropic effect of motion perception. If *α* = 0, *M*_*ij*_(*t, τ*) ignores the blind area of perception because the last two components in Eq.(2) always equal to 1 regardless of the relative positions of neighbors. Increasing *α* could make Eq.(2) amplify the anisotropic effect of motion perception, that is, the ability of individual perceiving movements of around neighbors gradually narrows to the front vision. Especially when *α* = 10, it induces *M*_*ij*_(*t, τ*) ≈ 0 when the neighbors’ relative positions are beyond the horizontal sight (−π/2, π/2) (**Fig. 2b**). Note that *M*_*ij*_(*T, τ*) = *M*_*ij*_(*T, τ*) if *α* = 0. Otherwise, *M*_*ij*_(*t, τ*) ≠ *M*_*ij*_(*t, τ*).

From the definition of *M*_*ij*_(*t, τ*) in Eq.(2), we derived that: (i) *M*_*ij*_is to evaluate the motion salience of neighbor-*j* from focal-*i*’s perception; (ii) MST indicates the duration of flying freedom of a neighbor; (iii) average MST reflects the degree of flying freedom of a flock. For example, Supplementary Fig. 13a-f show the trajectories, ⟨*M*_*ij*_⟩ (*τ*) and temporal velocity differences ***v***_i_(*t*) · ***v***_*j*_(*t*) of pair (1,2) and (2,4) from the mobbing flock shown in **Fig. 1a**. Pair (1,2) has larger MST due to bigger velocity difference, while pair (2,4) is totally opposite. Thus, for the whole flock of **Fig. 1a**, Supplementary Fig. 13g-i demonstrate that the MST and average velocity difference for all bird pairs display an obvious negative relation, that is, larger MST, bigger velocity difference.

#### Quantifying the leading tier of each individual from LF networks

The leading tier of each individual from an LF network of a flock within [*t* − *τ, t*] could be calculated by the local reaching centrality of an unweighted directed graph^37^ as following,

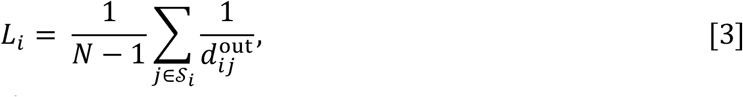

where *N* is the number of nodes, 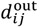 means the out-distance from node *i* to node *j*, and 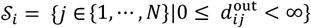 indicates the set of nodes with finite out-distance from node *i*. For example, in an unweighted directed graph, if node *i* without out-degree edges must locate at the bottom layer, then 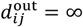 and *L*_*i*_ = 0; if node *i* with 1-step out-degree edges to the rest nodes must locate at the top layer, then 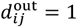 and *L*_*i*_ = 1. Therefore, *L*_*i*_ implies the hierarchical layer of each node in a directed network. Therefore, for a flock within the time period [*t* − *τ, t*], we first calculated the LF network, and consequently yielded the leading tier of each individual *L*_*i*_ (*t, τ*) by Eq.(3). Note that we set the non-zero elements in LF relation matrix as 1 when calculating *L*_*i*_ (*t, τ*).

#### The adaptive interaction based on perception of MS

To mimic the effects of MS observed in bird flocks, we introduced the adaptive MS-based interaction (AMS) to the classical self-propelled swarm model^23^. In the model, each agent moves towards a heading with a constant speed *v*_4_. The position of agent *i* is updated as

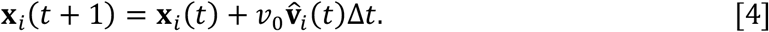

Hereinafter, 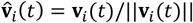 is the normalized velocity of agent *i*, and ***v***_*i*_(*t*) is updated as

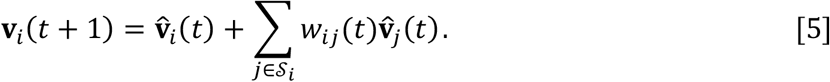

Here, 𝒮 _*i*_ represents the neighboring agents (except *i* itself) within a circle of sensing radius *r*_al_that is centered at agent *i* . *w*_*ij*_ is the weighted coefficient to reflect the heterogeneity of local interactions between individual pairs, which indicates the influences from neighbors exerted on the focal individual. According to the findings in bird flocks, AMS assumes that the neighbor with higher MS could impose much more influences on the focal individual. Thus, *w*_$%_, as the coefficient of adaptive interaction, could be quantified by MS in the form of,

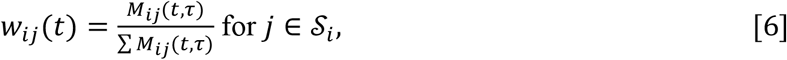

where *M*_*ij*_(*t, τ*) calculated by Eq.(2) equals the MS of neighbor *j* perceived by agent *i* within the period [*t* − *τ, t*].

Furthermore, to further investigate the effects of paying more attentions to neighbors with larger MS in AMS, we complemented a tunable percent parameter *x*% to select the neighbors from 𝒮 _*i*_ based on their MS values. For the focal agent *i*, (i) getting the metric-based neighbor set 𝒮 _*i*_ by the given distance criterion *r*_al_and calculating *w*_*ij*_ for *j* ∈ 𝒮 _*i*_ based on Eq.(6); (ii) picking up a part of neighbors as a new set 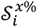 if the cumulative sum of *w*_*ij*_in a descending order is just larger than *x*%; (iii) renormalizing 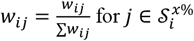 and setting 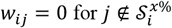. Here AMS-*x*% means at least the top *x*% of MS are involved in AMS. By this definition, the tunable *x*% could even further amplify the attention effect of neighbors with higher MS. For instance, if *x*% = 50%, AMS-50% makes the focal agent only align with those neighbors cumulatively possessing at least the top 50% MS and ignore the influences from neighbors with the latter 50% MS.

Unlike perceiving MS with a period of time as Eq.(2), there exists a kind of adaptive interaction according to transient heading difference (ATHD)^47^, such as,

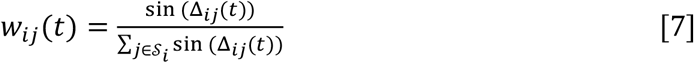

with Δ_*ij*_(*t*) = m*i*n {∣*θ*_*i*_ (*t*) – *θ*_*j*_ (*t*)o, 2π − ∣ *θ*_*i*_ (*t*) − *θ*_*j*_ (*t*) ∣ } ∈ [0, π] . *θ*_*i*_ (*t*) ∈ [−π, π] denotes the heading of agent-*i* at time *t*. This interaction strength adaptively increases with the increment of heading differences between neighbors and the focal individual. Once the difference is larger enough, i.e., larger than π/2 in Eq.(7), the contribution will decrease and eventually will become zero. Besides, *w* _*i* %_ = 1 for *j* ∈ 𝒮 _*i*_ makes the adaptive interaction degenerate to the standard Vicsek model^23^, which means the focal agent equally inteTracts with those neighbors located in the sensing radius.

In **Fig. 4**, we used a classical self-propelled particle model to compare the above different interaction rules as following^15,71^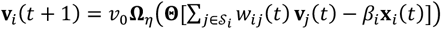 . In this model, all particles move at a constant speed *v*_4_ and align their directions of motion based on the local interaction rules, with some noise added. The operator Ω_F_ imposes the noise through rotating the vector by a random angle chosen from a uniform distribution with maximum amplitude of *η*, and operator Θ normalizes the argument to be a unit vector. Note that the term –*β*_*i*_x_*i*_ (*t*) is introduced as additional potential well that pushes particle *i* back towards the origin^71^. Here to mimic the leading effect, the potential well is only imposed on individual-1 (*β*) ≠ 0, red trajectories in **Fig. 4a-c**) to lead the flock come back to origin, and *β*_*i*_ = 0 for other particles. We ran this model in 3D without boundary conditions. Initially, particles are randomly distributed in a 3D sphere with a radius of *R* = 50m and moved in random directions with constant speed *v*_0_ = 10*m*/*s*. The parameters used in **Fig. 4** are *N* = 30, *r*_al_= 30*m*, Δ*t* = 0.05*s, η* = 0.05 and *β*_1_ = 0.08.We set the perceiving time *τ* = 20Δ*t* in AMS.

#### Swarm robotics system

To demonstrate the implication of AMS to enhance the self-organization of real swarming robots, we built a swarm robotics system that consists of ∼10^2^ magnitudes of miniature two-wheel differential mobile robots (Supplementary Fig. 17). Each robot is equipped with two stepper motors with reduction gears, a PCB board (with STM32F103RC8T6 microcontroller) for motion control and power management, a PCB board (with STM32F103RCT6 microcontroller and NRF24L01 wireless communication module) for communication and decision making, and a marker deck at the top for hosting passive infrared reflective balls (4-5 balls with diameter of 10mm) for localization (Supplementary Fig. 17a). The diameter of robot’s main body is 60mm and the diameter of marker deck at the top is 84mm. The robot, powered by two 3.7V rechargeable lithium batteries (2*800 mAh), can move according to specified linear and angular speed control commands. A NOKOV motion capture system is used to track the robot position x(*t*) (center of body) and heading *θ*(*t*), from which real-time linear speed *v*(*t*) and angular rate *ω*(*t*) of the robot are obtained by differential calculation. For simplicity, all motion control commands for swarming robots in our experiments are computed on a service computer and then simultaneously broadcasted to the swarm with a fixed time interval Δ*t* through a customized wireless communication protocol. After receiving the motion command of the desired linear speed *v*_*d*_(*t* + 1) and heading *θ*_*d*_ (*t* + 1) calculated from the swarm model, each robot performs the angular rate command *ω* = m*i*n*{*Δ*θ*/Δ*t, ω*_max_} within a control loop, where Δ*θ* = *θ*_*d*_(*t* + 1) − *θ*(*t*) and *ω*_max_is the maximum angular rate. To guarantee the transferability of the swarm model to hardware experiments, the system is capable of sending motion control commands as fast as 20Hz (i.e., support Δ*t* ≥ 0.05*s*) and capturing each robot’s motion states up to 300Hz. We took the maximum angular rate *ω*_max_ = 19.1 deg/s in the swarm experiments. See Supplementary Fig. 17c for the architecture of swarm robotics system.

Due to the limitation of arena size, we performed the swarm experiments with up to 50 robots (Supplementary Fig. 17b). However, to perform the swarm experiments with hundreds of robots, we transferred the real robots to semi-physical simulation with the same motion characteristic in Pybullet (Supplementary Fig. 20c and Supplementary Video 11).

## Data availability

All the experimental datasets analyzed in this study are either publicly available or kindly provided by the original authors.

## Code availability

All codes for real data analysis in this manuscript can be found at https://github.com/xiaoyandong08/Perception_of_Motion_Salience.

## Acknowledgements

We thank Professors Nicholas T. Ouellette and Dennis J. Evangelista for kindly sharing the flocking datasets. Y.D.X is supported by the National Natural Science Foundation of China (61902418). X.P. is supported by the National Natural Science Foundation of China (62076203).

## Contributions

Y.D.X. conceived and designed the project. Y.D.X. and X.P. managed the project. Y.D.X. performed all the real data analysis, analytical/numerical calculations and simulations. X.L. designed the collective following experiment. Y.D.X. designed the collective evacuation experiment. X.P., X.L., Z.Z., and Y.L.X. built the swarming robotic system. X.L., Y.L.X. and Z.Z. performed two swarm experiments. Y.D.X. wrote the manuscript. Y.-Y.L. edited the manuscript. All authors interpreted the results.

## Competing Interests

The authors declare no competing interests.

## Supplementary Materials

Supplementary Text

Figs. S1 to S25

Tables S1 to S2

Movies S1 to S11

## Supplementary Materials

Supplementary Text

Figs. S1 to S25

Tables S1 to S2

Movies S1 to S11

## Notes

### Competing Interest Statement

The authors have declared no competing interest.

